# Development and qualification of a high-yield recombinant human Erythropoietin biosimilar

**DOI:** 10.1101/2023.01.22.525046

**Authors:** Kakon Nag, Md. Jikrul Islam, Md. Maksudur Rahman Khan, Md. Mashfiqur Rahman Chowdhury, Md. Enamul Haq Sarker, Samir Kumar, Habiba Khan, Sourav Chakraborty, Rony Roy, Raton Roy, Md. Shamsul Kaunain Oli, Uttam Barman, Md. Emrul Hasan Bappi, Bipul Kumar Biswas, Mohammad Mohiuddin, Naznin Sultana

**Affiliations:** Globe Biotech Limited, 3/Ka (New) Tejgaon I/A, Dhaka 1208, Bangladesh

**Keywords:** Biologics, Cloning, Master cell bank, Bio-functionality, physiochemical, Characterization

## Abstract

Recombinant human erythropoietin (rhEPO) has been saving millions of lives worldwide as a potent and safe treatment for the lack of erythrocyte, which is caused by chronic kidney disease (CKD) and other issues. Several biosimilars of rhEPO have been approved since the expiry of the relevant patents to provide cost-effective options but the price of rhEPO is still high for the affordability of global community. Therefore, development of biosimilar of rhEPO at a lower price is highly necessary. Here we report the development and characterization of a biosimilar of rhEPO with high-yield satisfying regulatory requirements. The hEPO-expressing cDNA was stably expressed in CHO cells with successive transfection. The master cell bank (MCB) and working cell bank (WCB) were established from the best selected clone and characterized for 50 passages. The rhEPO was expressed from the WCB in single-use suspension culture system with a high-titer (1.24±0.16 g/L). To the best of our knowledge this is the highest reported rhEPO titer to date. The rhEPO was purified using a series of validated chromatography unit processes including virus inactivation and filtration. The purified EPO was formulated in serum-free buffer, sterile filtered, and analyzed as the biosimilar of reference product Eprex^®^. Physicochemical analysis strongly suggested similarities between the developed rhEPO (GBPD002) and the reference. The *in vitro* and *in vivo* functional assays confirmed the similar biofunctionality of the GBPD002 and Eprex^®^. GBPD002 could provide a less-expensive solution to the needful communities as an effective and safe biosimilar where rhEPO treatment is necessary.

## 1. INTRODUCTION

Erythropoietin (EPO) is an essential hormone for erythrocyte or red blood cell (RBC) production in mammals [1]. It is also known as haematopoietin or hemopoietin. EPO is a glycoprotein and composed of 165 amino acids with an estimated molecular weight of 34 kDa [2]. It has multiple isomeric forms; 5 – 8 isoforms were identified over the isoelectric point (pI) range of 4.4 – 5.2 [3]. The 3 potential N-linked glycosylation sites were predicted located at Asn24, Asn38, Asn83, whereas the single O-glycosylation site was predicted on Ser126 [4-7]. Approximately, 40% molecular weight of the protein has been attributed to the glycosylation. EPO has two disulfide bonds as well, which are formed between the cysteines of 7-161 and 29-33 amino acids; these cysteine-cysteine bridges are essential for maintaining biological activity [2, 8].

EPO is secreted from the kidney, which is enhanced in response to cellular hypoxia [9]. It promotes the division and differentiation of bone marrow resident committed erythroid progenitors to maintain erythrocyte population in blood for oxygen supply [10]. EPO is the primary erythropoietic factor that cooperates with various other growth factors like, interleukin (IL)-3, IL-6, glucocorticoids, and stem cell factors (SCF), which are involved in the development of erythroid lineage from multipotent progenitors. EPO works on RBC-progenitor by binding to the EPO receptor located on the surface of these cells, which triggers JAK2 signaling cascade [11, 12]. Activated JAK2 then initiates the STAT5, PIK3 and Ras MAPK signaling cascades resulting in differentiation, survival and proliferation of the erythroid cell [13].

Compromised EPO production is the main reason of anemia in human with chronic renal failure [14]. It has been demonstrated that the human recombinant epoetin (rhEPO) stimulate erythropoiesis in anemic patients with chronic kidney disease (CKD), including patients who need and do not need dialysis [15-18]. The administration of rhEPO has been found beneficial for the treatment of chemotherapy-induced anemia in cancer patients and to reduce the requirement for allogenic blood transfusions in patients with mild anemia who are undergoing surgery [19-21]. Furthermore, rhEPO is recommended for patients who are at high risk for perioperative transfusions due to substantial blood loss. Though, the use of rhEPO was initially restricted to dialysis patients with most severe forms of anemia, however, it has been currently administered to most dialysis patients with renal anemia and to predialysis patients [22].

The entry of rhEPO into clinical practice was a milestone for the treatment of renal anemia [23], followed by alfa, beta, zeta and other formulations. Johnson & Johnson’s Eprex®/Procrit was the first rhEPO preparation that was approved by the US FDA and the European Medicine Agency (EMA) in June 1989; Amgen’s Epogen was followed later with the approval of US FDA in February 1999 [24, 25]. The patents on Epogen/ Eprex^®^ already expired in both the US and in Europe in 2013 [26], and opened up the opportunity for developing biosimilars for rhEPO to supply cost-effective medication to the market. Three biosimilars of rhEPO, *viz*., Binocrit® from Sandoz, Abseamed® from Medice and Epoetin Alfa Hexal® from Hexal AG were approved by the EMA, and Retacrit (epoetin alfa-epbx) from Hospira (Pfizer) was approved in the US in May 2018 [24].

More than 850 million people worldwide were estimated who have been suffering from CKD [24]. The prevalence of anemia in patients with CKD (15.4%) in the USA is double compared with the overall population (7.6%); the prevalence rate increased to 53.4% with stage 5 of the disease [27]. CKD patients with anemia are at a higher risk of health-related quality of life impairment, cardiovascular disease, end-stage renal disease, hospitalization and mortality, compared with CKD patients without anemia [1, 28, 29]. The high prevalence of CKD-related anemia has been creating growing demand for the rhEPO but limited supply of qualified rhEPO has made the treatment expensive. The cost to treat anemia by rhEPO per quality-adjusted life-year was estimated at US $24,128.03 and US $28,022.33 for the Hb level 9-10g/dl and 11-12g/dl, respectively [30]. The average GDP per capita of the global population, which was US $11570 in 2019 [31], is below the average medication cost of EPO treatment.

Biosimilar is the fastest-growing segment in the biological drug market, owing to the value points of comparatively low-cost therapeutics than the originator with desired efficacy [25, 32]. Despite introduction of few biosimilars in the market [26], the price of rhEPO is still out of reach for global community, and inviting the introduction of more cost-effective and affordable biosimilars to the world. Therefore, we have developed a biosimilar for rhEPO with the objective to reduce the cost by exploiting the science and technology. Here, we describe the development and characterization of GBPD002, a candidate biosimilar of rhEPO, with a superior productivity that will help to cater the product for better socioeconomic benefits to the global community.

## 2. MATERIALS AND METHODS

### Amplification of the gene of interest

Human spleen total RNA (Thermo Fisher Scientific, USA) was used as template for cDNA synthesis using Superscript IV 1^st^ strand (Thermo Fisher Scientific, USA) cDNA synthesis kit. Primers were designed against EPO amino acid sequence obtained from DrugBank (accession number DB00016) and nucleic acid sequence obtained from GenBank (accession number X02158.1), for the cDNA synthesis and subsequent amplification of the gene of interest. Platinum Pfx DNA polymerase (Thermo Fisher Scientific, USA) was used for the amplification using standard PCR protocol (Denaturation: 94 °C for 30 secs, annealing: 66 °C for 30 secs, extension: 68 °C for 42 secs and total number of cycles: 35) in ProFlex 3×32-well PCR system (Applied Biosystems, USA). Desired amplified product was excised from 1.2% agarose gel and purified using GeneJET gel extraction and DNA cleanup kit following supplier’s protocol (Thermo Fisher Scientific, USA).

### Cloning

pcDNA5/FRT (Thermo Fisher Scientific, USA) mammalian expression vector was used for construction of the rDNA. Amplified gene of interest and expression vector was digested using FastDigest XhoI and HindIII (Thermo Fisher Scientific, USA) restriction endonuclease following standard protocol. The digested sticky end insert and vector was purified from agarose gel excision using GeneJET gel extraction and DNA cleanup kit. The insert was then ligated with the pcDNA5/FRT vector using T4 DNA ligase (Thermo Fisher Scientific, USA) following standard sticky end ligation protocol. After ligation, the constructed rDNA was transformed into Top10 chemically competent *E. coli* cells (Thermo Fisher Scientific, USA) for rDNA amplification. The rDNA purification was done using PureLink quick mini and PureLink plasmid midi kit (Thermo Fisher Scientific, USA) following supplier’s protocol for mini and midi scale of rDNA, respectively.

### Digestion and Sequencing

XhoI and HindIII was used to digest the gene of interest from the purified rDNA and insert size was confirmed by agarose gel electrophoresis. BigDye Terminator v1.1 Cycle sequencing kit (Thermo Fisher Scientific, USA) was used for sequence amplification, 3500 genetic analyzer (Applied Biosystems, USA) was used for sequencing and analysis was done using ABI Sequencing Analysis v5.4 software.

### Cell transfection

Flp-In system (Thermo Fisher Scientific, USA) was used for stable transfection. Flp-In CHO cells were chemically co-transfected with the linearized rDNA and circular pOG44 Flp-recombinase vector using Lipofectamine 3000 (Thermo Fisher Scientific, USA) following their protocol. Constructed rDNA was linearized using FastDigest LguI (SapI) (Thermo Fisher Scientific, USA). Approximately, 0.75×10^5^, 1×10^5^ and 1.5×10^5^ cells in FBS containing Ham’s F-12 nutrient media were seeded in one 24-well TC-treated sterile cell culture plate (SPL, Korea) 24 hours before transfection. Wells seeded with 1×10^5^ and 1.5×10^5^ cells reached at desired 60 to 70% confluency with a viability of around 90% and transfected with 2.5 µg linear rDNA and 22.5 µg pOG44 in Opti-MEM media. After 6 hours of transfection, the cells were transferred in Ham’s F-12 media for protein expression. Another expression construct was made using the pLVX-EF1a/puromycin (+) plasmid (Genemedi, China) containing EF1a promoter. This plasmid also includes puromycin resistance gene and the ampicillin resistance gene as known selectable markers of stable mammalian transfectants. This plasmid was used for second transfection to obtain high-copy number of insertions in the target cells.

### Antibiotic selection

20,000 cells per well were seeded in a 24-well TC treated plate for applying the selection pressure. Hygromycin B (Thermo Fisher Scientific, USA) of different concentrations (200 µg/mL to 1000 µg/mL, with 100 µg/mL gradual increment) was used in 3 replicate wells for each concentration with negative control 24 hours post seeding. Antibiotic containing media of respective concentration was changed in every 48 hours for 14 days. The selection pressure was withdrawn from the wells on day 14, where 600 µg/mL of Hygromycin B was maintained and the cells were split into a fresh 24-well plate. Western blot analyses of all representative wells were performed to check the expression of desired protein. After second transfection, different concentration of puromycin antibiotic (25 µg/mL to 200 µg/mL, with 25 µg/mL gradual increment) challenge was applied. After that 100 µg/mL of puromycin was maintained and the cells were split into a fresh 24-well plate.

### Clonal isolation and stable expression analysis

After antibiotic selection and expression analysis, clonal isolation following serial dilution method was performed in a 96-well plate, with a starting cell of 10,000 at A1 well. Serial dilution was done in 10% FBS containing Ham’s F12 media in treated plate. After that, western blot was performed for expression analysis. Four top performing clones were selected and were subjected for re-transfection followed by clonal isolation. For expansion, cells were cultured in 75 cm^2^ TC-treated flask. The stable expression of the desired proteins was periodically checked by Western blotting using the media in culture [32, 33]. After second antibiotic selection, clonal isolation was performed by automated cell shorter BD FACSAria™ Fusion Flow Cytometers (BD Bioscience, USA). After that, western blot was performed and top eight clone were selected.

### Best clone selection and MCB development

Four clones (Clone No. GB002EC001, GB002EC002, and GB002EC003) were processed forward. These clones were systematically adapted to suspension culture in serum free medium. The serum-free suspended cells were cultured in microbioreactor (Ambr® 15 cell culture bioreactor; Sartorius, Germany) to determine the specific productivity (*Q*_p_) and cellular growth rate of clones. Based on the productivity and process suitability the best clone was selected. Cell viability, growth rate, specific expression kinetics, cell morphology, stability of the desired protein in culture etc. were considered for selection. The dot blot and Western blot analyses were performed routinely for evaluation. The stability of the selected clone was confirmed up to 50^th^ passages (generally, 4 days/passage) over a period of 200 days.

### Cell bank identity

The GBPD002 (rhEPO) expressing suspended CHO (expressing clone GB002EC003) cells were seeded into 24-well TC-nontreated plate containing 600 µL of 100% ActiPro medium and four replicates was prepared for seeding density of approximately 4.80E+04 cells/600pl or 3.0E+05 cells/mL. Cells were counted every 24 hours for up to day 18, with the daily replenishment of media. The media was collected and analyzed for the expression of desired protein by Western blot. Growth rate (p) and population doubling time (PDT) is calculated from the projected graph. Cell morphology and cell viability analysis were done by microscopic evaluation of MCB. The growth kinetics of the cells were measured over the 50 passages.

### Dot blot analysis

Dot blot analysis was performed to identify the best clone. PVDF membrane (Thermo Fisher Scientific, USA) was taken and grid was drawn by pencil to indicate the region of blot. To activate the membrane, it was submerged into 100% methyl alcohol for 3 minutes at room temperature. Then the membrane was incubated in transfer buffer (pH 8.3) for 12 minutes. After proper incubation the membrane was taken on a wet Whatman filter paper. 10 µL sample from each clone was added on each grid of the membrane. Then the membrane was dried at 40 °C for 5 minutes and activated the membrane again by incubating it into 100% methyl alcohol for three minutes at room temperature without any shaking. Then membrane was transferred into transfer buffer (pH 8.3) for 12 minutes. After that, the membrane was blocked with blocking buffer for 1 hour at room temperature. After successful blocking, 1 µL EPO specific primary antibody (Thermo Fisher Scientific, USA) was added with 10 mL blocking buffer for the preparation of primary antibody solution and membrane was submerged into it for 1 hour at room temperature. The membrane was washed with TBST buffer for 10 minutes. This process was performed three times. After that, 1 µL goat anti rabbit secondary antibody (Thermo Fisher Scientific, USA) was added with 10 mL blocking buffer for the preparation of secondary antibody solution and membrane was submerged into it for 1 hour at room temperature. Again, membrane was washed with TBST buffer for 10 minute and performed this step three times. Then, membrane was rinsed with type-1 waster for two minutes and placed into a clean plastic sheet. Novex® ECL chemiluminescent substrate (Thermo Fisher Scientific, USA) was added over the entire blotting paper and incubated for two minutes and finally image was captured by Amersham Imager 600 RGB [34].

### SDS-PAGE analysis

SDS-PAGE was performed under reducing conditions. In short, the samples and the reference product Eprex^®^ [@6.5 μL (1 µg) in 2.5 μl NuPAGE^®^ LDS Sample Buffer (4×) and 1 μl NuPAGE^®^ Sample Reducing Agent (10×)] were loaded in Bolt 4-12% Bis-Tris Plus Gels (Thermo Fisher Scientific, USA). Samples were denatured at 95°C for 3 mins, and cold on ice, and spin them down (for 1min, at 12,000 rpm) before loading them in gel. Proteins were separated using 90 mA for 45 min. Proteins were visualized by Coomassie brilliant blue staining. Novex sharp pre-stained protein marker (Thermo Fisher Scientific, USA) was used as reference for molecular weight.

### Western blot

Comparative analysis was done between reference product Eprex^®^ and GBPD002 protein sample using Western blot at reducing conditions. Bolt 4-12% Bis-Tris Plus Gels (Thermo Fisher Scientific, USA) was used for separation of the proteins by molecular weight and then separated proteins were transferred in iBot 2 PVDF membrane (Thermo Fisher Scientific, USA). Anti-Epo polyclonal antibody (Thermo Fisher Scientific, USA) was used as a primary antibody and HRP conjugated Goat anti-rabbit (H+L) IgG (Thermo Fisher Scientific, USA) was used as a secondary antibody for detecting target proteins. Molecular size was determined by using novex sharp pre-stained protein marker (Thermo Fisher Scientific, USA). Novex^®^ ECL Chemiluminescent Substrate (Thermo Fisher Scientific, USA) for HRP was used to detect the signal and the imaging was done using Amersham Imager 600 RGB (GE Healthcare, USA).

### Two-dimensional gel electrophoresis

Two-dimensional (2D) gel electrophoresis was performed using 20 μg of each sample for charge variant analysis. In short, isoelectric focusing (IEF) was performed in ZOOM® Strip pH 3–10L (Thermo Fisher Scientific, USA) followed by SDS PAGE in Novex™ 4-¬20% Tris-¬Glycine ZOOM™ Protein Gels (Thermo Fisher Scientific, USA) with XCell SureLock™ Mini-Cell Electrophoresis System (Thermo Fisher Scientific, USA). Gels were stained with Coomassie blue and evaluated against Novex sharp pre-stained protein marker (Thermo Fisher Scientific, USA). The imaging was done using Amersham Imager 600 RGB (GE Healthcare, USA).

### Insert sequence and the copy number of insert

The DNA sequencing of inserted EPO gene in MCB and WCB were performed as described elsewhere in the article using the Chromosomal DNA and cDNA. The copy number of inserts in MCB and WCB were analyzed using QuantStudio 12K Flex system qRT-PCR system (Thermo Fisher Scientific, USA) and genomic DNA from sample cells. Comparative C_t_ and melt-curve analysis method was used to identify the relative copy number of inserts in relevant genome. A Flp-In-CHO stably transfected with rhEPO is considered as the positive reference for relative quantitation of the copy number of the inserted rhEPO DNA in WBC. A serial dilution of the reference cells was considered as scale for calculating copy numbers of the experimental sample.

### Mycoplasma detection

Comparative C_t_ and melt-curve analysis were done for mycoplasma contamination check for MCB and WCB. Analysis was done by using MycoSEQ™ Mycoplasma Real-Time PCR Detection Kit (Thermo Fisher Scientific, USA) and QuantStudio 12K Flex system qRT-PCR system (Thermo Fisher Scientific, USA). Positive control reaction, negative control reaction, inhibition-control reaction and extraction spike control were used for method suitability test.

### *Upstream* processing

Batches were started from WCB at an N-4 step. Three consecutives’ batches were performed in 5 L bioreactor using ReadyToProcess WAVE 25 system (GE Healthcare, USA). A single-use 10 L Cell bag™ bioreactor (working volume: 0.5-5 L; GE Healthcare, USA) was placed on the rocker. Cell cultures were controlled using the UNICORN™ system control software (GE Healthcare, USA). Cells were seeded at ∼ 7.93 × 10^8^ total viable cells (around 7.93 × 10^5^ viable cells/mL) in a chemically defined media (ActiPro media; GE Healthcare, USA). The cultures were maintained with setpoints for pH at 7.2±0.05, dissolved oxygen (DO) at 40% air saturation and stage-specific variable rocking speeds from 18 – 28 with rocking angle 5 – 8 degrees. Culture pH was controlled by either addition of 7.5% NaHCO_3_ and/or by sparging of CO_2_ gas. Bioreactor cultures were provided with gas flow from 0.10 – 0.5 lpm through air sparger, and DO was controlled by sparging with air and oxygen. Culture temperature was kept at 37 °C from day 0 to day 7, and shifted to 32 °C on the day 7. Required amounts of media and nutrients were added to the cultures on regular basis. The cultures were harvested at day 17. Samples were taken at every 24-hour interval for measurement of viable cell density (VCD), viability, and productivity. The media were harvested in 2D sterile bags after filtration through 0.6 µm followed by 0.2 µm PES filter (GE Healthcare, USA) and proceeded for downstream processing.

### Downstream processing & formulation

The binding and elution conditions in each of chromatography processes were screened out in 96 deep-well plate format and adapted in small scale column format. The optimum conditions (volumetric flow rate, linear flow rate, residence time, dynamic binding capacity of each resin, operating temperature etc.) for each chromatography step were determined in small scale. The unit processes were scaled up, and the conductivity and pH of samples were considered as in-process check (IPC). The rhEPO was enriched onto 0.40 L Capto blue (GE Healthcare, USA) column with AKTA Avant 150 system (GE Healthcare, USA) followed by washing the unbound potential contaminants, and then eluted using gradient of 20 mM Tris-HCl, pH 7.4 and 1.5 M NaCl. FPLC systems and columns were sanitized with 1.0 N NaOH prior and after each purification step. The in-line UV_280nm_, conductivity and pH of the sample were considered as IPC. The eluate from Capto blue was diafiltered against Tris-HCl, pH 7.0 to remove the salt and undesired low molecular weight (LMW) molecules (<10 kDa) from sample using 0.10 m^2^ Sartocon slice PES cassette (Sartorius Stedim, Germany) and AKTA flux 6 system (GE Healthcare, USA). The system and cassette were sanitized with 1.0 N NaOH before and after diafiltration. The conductivity and pH of retentate were considered as IPC. The retentate was loaded onto 0.25 L Q Sepharose HP (GE Healthcare, USA) column to bind desired isoforms. The rhEPO was separated from some host proteins (HCPs), nucleic acid (DNA) and adventitious viruses using gradient of Tris-HCl, pH 7.0 and 1.5 M NaCl. The in-line UV_280nm_, conductivity and pH of the sample were considered as IPC. The elute from Q Sepharose was loaded on to 0.20 L source 15RPC (GE Healthcare, USA) column to separate non-glycosylated and glycosylated rhEPO. The eluate was collected using gradient of 0.1% TFA in WFI and 95% acetonitrile. The in-line UV_280nm_, pH and conductance of the eluate were considered as IPC.

The eluate of source 15RPC was incubated for virus inactivation at low pH. The virus-inactivated RPC eluate was loaded on to 0.10 L Macrocap SP Sepharose (GE Healthcare, USA) column to remove RPC buffers and rhEPO aggregates followed by reconstitution into pre-formulation buffer. After washing with 20 mM glycine buffer, the sample was eluted using gradient of 25 mM sodium phosphate, 0.00023 mM polysorbate 80, pH 7.0 and 1.0 M NaCl. The in-line UV_280nm_, conductance and pH of the eluate were considered as IPC. The eluate of Macrocap SP was filtered through 20 nm Virosart HF (Sartorius Stedim, Germany). The rhEPO concentration was checked and diluted to desired concentration in formulation buffer. After sterile filtration through 0.45|0.2-micron PES Sartopore 2 filter (Sartorius Stedim, Germany) the formulated bulk sample was collected in 3 L Flexboy 2D bag (Sartorius Stedim, France) and filled into pre-sterile syringe (Schott, Switzerland) for dose preparation. The pre-filled syringes were stored at 5 ± 3 °C and were subjected for analysis as per specification.

### Enzyme-linked immunoassay (ELISA)

Microtiter plates (Nalge Nunc International, USA) were coated with polyclonal anti-human EPO (4 mg/mL; Thermo Fisher Scientific, USA) in 0.1 M sodium bicarbonate buffer (pH 8.3) at 4°C overnight. The plates were blocked by 3% BSA/PBS for 2 h and then incubated with serial dilutions of an EPO standard (Thermo Fisher Scientific, USA)) or culture supernatant samples for 4 hours at room temperature. After washing, the bound EPO was incubated with monoclonal mouse anti-EPO antibody (1 mg/ mL; Thermo Fisher Scientific, USA) at 4°C overnight and then alkaline phosphatase (AP) conjugated anti-mouse IgG adsorbed with rat serum protein) (Thermo Fisher Scientific, USA) for 2 h diluted 1: 15,000 in 1% BSA/PBS/0.05% Tween20. For the detection of the antigen–antibody reaction, p-nitrophenyl phosphate was added as a substrate, and incubated at 4°C overnight. The optical absorbance at 405 nm was measured by an ELISA reader. Each incubation step was followed by washing four times with PBS/0.05% Tween20.

### Chromatography for determination of assay and impurities

The quantitative analysis (assay) for rhEPO of the samples were performed using a Vanquish UHPLC system (Thermo Fisher Scientific, USA). The reference product Eprex^®^ and samples (20 µL of each) were applied in RPC column (Hypersil GOLD C8, 175 Å, 2.1× 100 mm, 1.9 µm column; Thermo Fisher Scientific, USA) using an auto injector system. A gradient of media A (water with 0.1% FA) and media B (90% ACN in water with 0.1% FA) was used as mobile phase at a flowrate of 0.3 mL/min. The formulation buffer was considered as reference for base line signal. The impurity profile of the samples was analyzed using size exclusion chromatography (SEC). A 50 µL of each sample and reference product Eprex^®^ were loaded in a Biobasic SEC-300 column (300 mm*7.8mm, 5µm; Thermo Fisher Scientific, USA), and the analyses were performed using an Ultimate 3000 RSLC system (Thermo Fisher Scientific, USA). Phosphate buffer with pH 7.4 was used as carrier phase at a flow rate of 1.0 ml/min. All analyses were performed with a 25 min run time and at a fixed column temperature of 70 °C. The chromatograms were obtained from the absorbance at 280 nm, and relevant values were used for calculations.

### Peptide mapping

40 µg of rhEPO preparation was diluted in 50 mM ammonium bicarbonate (Wako Pure Chemicals Industries Ltd., Japan), at pH 8 containing 8 M urea (Thermo Fisher Scientific, USA). To reduce the sample, 500 mM DTT (Thermo Fisher Scientific, USA) was added to the solution to a final concentration of 20 mM (1:25 dilution), mixed briefly, and then incubated at 60 °C for 1 hour. For alkylation, 1 M IAA (Sigma-Aldrich, USA) was added to the reduced protein sample to a final concentration of 40 mM (1:25 dilution), and incubated the reaction mixture for 30 minutes protected from light. The reaction was stopped by adding 500 mM DTT solution to a final concentration of 10 mM (1:50 dilution). Trypsin (Thermo Fisher Scientific, USA) was added to the sample solution to a final trypsin to protein ratio of 1:23 (w/w). Samples were incubated at 37 °C for 16 – 24 hours. Reaction was stopped by adding formic acid up to bringing the pH 2.0. C18 spin column (Thermo Fisher Scientific, USA) was prepared as per manufacturer’s instruction to purify the peptide-pool. Columns were activated by adding 200 µL 50% acetonitrile (Wako Pure Chemicals Industries Ltd., Japan), and equilibrated using 200 µL of 0.5% formic acid (Wako Pure Chemicals Industries Ltd., Japan) in 5% acetonitrile (Wako Pure Chemicals Industries Ltd., Japan). Samples were applied to the column and eluted using 20 µL of 70% acetonitrile. Samples were dried under low temperature and vacuum and processed for experiments.

rhEPO preparation was digested with Serine Protease (MS grade, Pierce, Thermo Fisher, USA) and purified according to the supplier’s instructions. 2 µg of digested peptides were loaded into mass spectrometry system (Q Exactive Hybrid Quadrupole-Orbitrap MS, Thermo Fisher Scientific, USA). Hypersil gold C18 (100×2.1 mm; particle size: 1.9 µm, Thermo Fisher Scientific, USA) column was used for separation of peptides. Column oven temperature was set at 40 °C and samples were eluted in 95– 60 % mobile phase A (0.1% formic acid in water) and 5 – 40 % mobile phase B (0.1 % formic acid in acetonitrile) gradient with 0.300 mL/min flow rate for 100 minutes. Peptide elution were checked by absorbance at 214 nm. For peptide identification, data-dependent mass spectrometry was performed where full-MS scan range was 350 m/z to 2200 m/z, resolution was 70000, AGC target was 3E6, maximum IT was 100 milliseconds (ms). Data-dependent mass spectrometry resolution was 17500, AGC target was 1E5, and maximum IT was 100 ms. Data analysis was performed in BioPharma Finder (Thermo Fisher Scientific, USA) using variable parameters to get confident data, and then data were combined in one map to visualize complete fragmentation.

### Glycosylation pattern analysis

20µg of rhEPO and reference product Eprex^®^ were treated with 20 units of recombinant PNGase F (Thermo Fisher Scientific, USA) at 37 ºC for hours. Samples were subjected for SDS PAGE and Western blot analyses as described elsewhere in the article. The imaging was done using Amersham Imager 600 RGB (GE Healthcare, USA).

### Host Cell DNA

Host cell DNA were analyzed using resSEQ™ CHO DNA Quantification Kit (Thermo Fisher Scientific, USA) following supplier’s protocol. Briefly, six 10-fold serial dilutions of control CHO DNA were prepared using DNA dilution buffer such that the final amount of CHO DNA in each reaction ranged from 3000 – 0.03 pg. Forward and reverse primers and a FAM-labeled probe that were designed to amplify a hamster-specific region of a multi-copy gene were provided. Each 30-µl PCR reaction mix contained 2 µl of negative control, 3 µl of 10x primer/probe mix, 15 µl of 2x Environmental Mastermix, and 10 µl of diluted CHO DNA. All reactions were performed on the QuantStudio 12K Flex System (Thermo Fisher Scientific, USA) using the following cycling conditions: 10 minutes at 95 °C to activate the enzyme, followed by 40 cycles of 15 seconds at 95 °C and then 1 minute at 60 °C.

### Host Cell Protein

Host cell Protein were analyzed using ProteinSEQ™ CHO HCP Quantitation Kit (Thermo Fisher Scientific, USA) as per supplier’s protocol. Briefly, seven 5-fold serial dilutions of control CHO HCP standard were prepared using DNA dilution buffer such that the final amount of HCP standard in each reaction ranged from 3125 – 0.2 ng/mL. Then, MagMAX express-96 magnetic particle processor (Thermo Fisher Scientific, USA) was used for capture the HCP automatically followed by several steps; i) preparing wash plate ii) preparing qPCR plate, iii) preparing probes plate, iv) preparing capture plate. Finally, QuantStudio 12K Flex System (Thermo Fisher Scientific, USA) using the following cycling conditions: 10 minutes at 37 °C to activate the enzyme, 20 seconds at 95 °C followed by 40 cycles of 15 seconds at 95 °C and then 30 seconds at 60 °C.

### Particle size distribution

To stabilize the system, the equipment was turned on minimum 30 – 60 minutes before taking the reading. Samples were prepared in 1× PBS (pH 7.2) and passed through 0.22-micron filter. Respective buffers were used as dispersant. The samples were allowed for stabilization at 20 ºC for 20 min, and then analyzed in disposable plastic cuvette using a Zetasizer Nano ZSP (Malvern Panalytical Ltd., UK). The refractive index (RI), viscosity and dielectric constant of dispersion buffer (1× PBS, pH 7.2 at 20 °C) was considered 1.33, 0.88 cPs and 79, respectively.

### Receptor binding

The Biacore T200 equipment (GE Healthcare, USA) was used for relevant experiments. EPO receptor protein (Fc chimera active; Abcam, USA) was immobilized on Series S Sensor Chips CM5 (GE Healthcare, USA) using amine coupling kit (GE Healthcare, USA). First, the flow-cell surface of Series S Sensor Chips CM5 was activated by injecting a mixture of EDC/NHS (1:1) for 7 minutes. Then 70 µL of 50 µg/mL EPO receptor protein was prepared in sodium acetate at pH 5.0 and injected over the activated surface at 10 µL/min flow rate. Residual NHS-esters were deactivated by a 70 µL injection of 1 M ethanolamine, pH 8.5.

The immobilization procedure was performed in running buffer HBS-EP, pH 7.4 (GE Healthcare, USA). rhEPO samples were passed over the active flow cell surface of CM5 chip for kinetics analysis. Glycine-HCl of pH 2.5 was used for regeneration. All samples were diluted in 1 x HBS-EP running buffer (pH 7.4).

### In vitro functional assay using cell culture

TF-1 cell line, which was originally derived from a patient diagnosed with erythroleukemia, were used for the *in vitro* functional assay of rhEPO. Cells were maintained in RPMI 1640 medium (Gibco, USA) supplemented with 10 % FBS (Thermo Fisher Scientific, USA), 1% PS (Thermo Fisher Scientific, USA), and 5 ng/mL human recombinant GM-CSF (Thermo Fisher Scientific, USA). The incubator condition was set at 37 °C with 5 % carbon dioxide environment. During the experiment, GM-CSF were eliminated from the cells by washing twice in PBS. Cells were then seeded at a density of 10^5^ cells/well in 24-well TC-nontreated cell-culture plate. Cells were allowed to grow for 72 h in the presence or absence either rhEPO samples at indicated concentrations. Eprex^®^ at indicated concentrations were used as positive control. Cells were collected after 72 hours through centrifugation for 10 min at 800 ×*g*. After washing with PBS, cells were counted using an automated cell counter (Countess 2, Thermo Fisher Scientific, USA)

### In-vivo assay by mice model

64 albino normocythemic mice (8-12 weeks old) were randomly assigned to one of eight the (groups) cages at day (-)2. The groups were as follows: control, placebo (formulation buffer), treatment I (20IU/mL, GBPD002), treatment II (40IU/mL, GBPD002), treatment III (80IU/mL, GBPD002), treatment I (20 IU/mL, reference), treatment II (40 IU/mL, reference), and treatment III (80 IU/mL, reference). There were eight mice in each cage. Animals from each group received a pre-defined 0.05 mL subcutaneous injection at day 0. After the injections, on day 4, the blood samples were collected from the animals. RBC and Hemoglobin were analyzed using auto hematology analyzer (BK-6190 VET, Biobase, China). For reticulocyte analysis, blood samples were diluted 500 times in the buffer and thiazole orange (Sigma-Aldrich, USA) was added. After staining for 3–10 min, cells are washed and the reticulocyte count was analyzed using a flow cytometer (BD FACSLyric™ Flow Cytometer, BD Bioscience, USA).

### Sterility and endotoxin testing

Bacterial and fungal sterility were tested using direct inoculation technique. Samples (1 mL) were inoculated in 10 mL of Tryptic Soya Broth (TSB) media for 14 days and absorbance were measured at 600 nm. Endotoxin in samples were tested using Pierce LAL Chromogenic Endotoxin Quantitation Kit (Thermo Fisher, USA) as per supplier’s instructions.

## 3. RESULTS

Development and characterization of a biosimilar is a meticulous process that has to be aligned with the regulatory framework. A snap shot of the overall workflow for the study is shown in figure-1. The activities mentioned in the article have gone through according to the work breakdown structure (WBS) of (Fig.: 1A), which is designed to comply regulatory requirements for a biosimilar.

**Figure 1:**
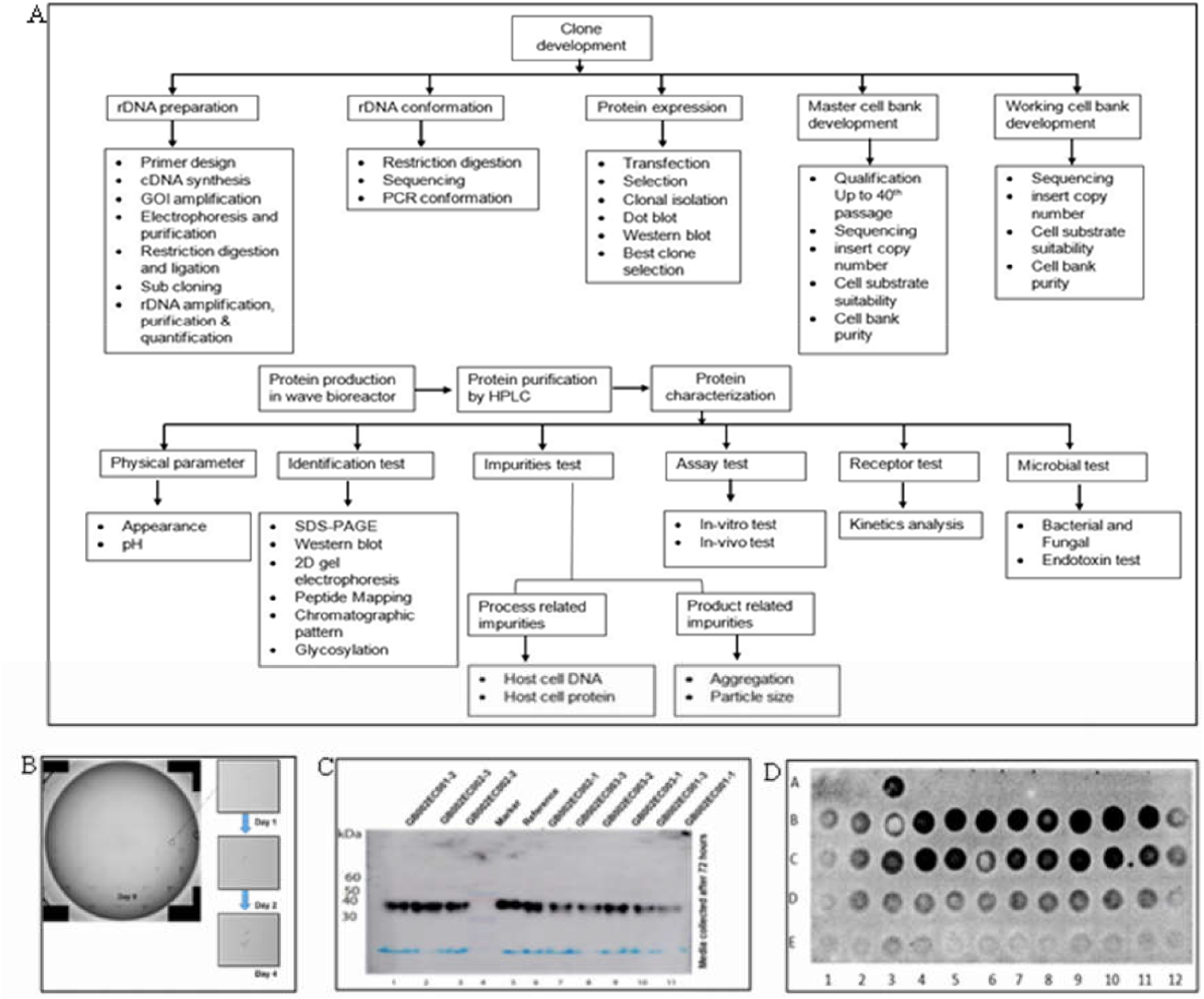
(A) Work break down structure of development and qualification of GBPD002 (B) Clonal isolation, (B) Western blot analysis of isolated expression clones. (C) Dot blot analysis for protein expression to select the best clone.

### rhEPO expression construct and high-level rhEPO-expressing cell line development

The amplified cDNA of rhEPO from human spleen was cloned into pcDNA5/FRT plasmid. Through a tBLASTn search against human genome, the DNA sequence of the insert data was confirmed to have an amplicon of 611 nucleotide (nt) with the desired 501 nt-long ORF of hEPO with start and stop codon (accession no.: X02158). The coding sequence was found 100% match to translate the hEPO protein (DrugBank reference no.: DB00016) with the pre-sequence. After transfection of the expression plasmid in Flp-In-CHO cells followed by challenge against Hygromycin B at 600 µg/mL concentration for 14 days, we have observed the existence of high-level of rhEPO-expressing cells. Lower concentration of Hygromycin B challenges was associated with low-level of rhEPO expression. We have screened the cells against 800 µg/mL of Hygromycin but no cells were survived. Several single clones were identified (Fig.: 1B) as rhEPO-expressing cells from the surviving pool of 600 µg/mL Hygromycin B challenge. They were amplified in separate well and the expression of the rhEPO was confirmed by Western blot. Most of them were found equipotent in terms of the rhEPO expression level (Fig.: 1C). After second transfection on these cells, we were able to identify surviving cells even in 100 µg/mL puromycin challenge, which represents higher level of antibiotic resistivity in these cells. Several single colonies were identified and allowed to grow. After dot blot analysis we found several of these clones were having significant higher level of rhEPO expression (Fig.: 1D).

We have selected eight of the similar high-level expressing cells with desired cellular morphology of a round shape isolated single cell in suspension from this set of clones. On a culture system with daily media replenishment, we found that four of the clones were growing above the other cells suggesting that they have higher expression level and higher cell growth kinetics as well. These cells were tested again for expression level of rhEPO with daily media replenishment and cell population correction to have the similar cells altogether throughout the experiment period of 7 days. We found that the relative specific expression of rhEPO for these cells were more or less similar (Clone no. 1: 5.4 pg/cell/day; Clone no. 2: 5.8 pg/cell/day; Clone no. 3: 5.5 pg/cell/day; and Clone no. 4: 5.7 pg/cell/ day. Since these clones have similar growth kinetics and expression profile therefore, they were further subjected for stress-screening mimicking mid- to end-stage fed-batch environment. These cells were allowed to grow in fixed media (no media change over 7 days) to check the tolerance capacity of the individual clones against media deprivation and media toxicity due to the continuous accumulation of cellular metabolic load. We found that though they have similar morphology, growth kinetics in fresh media and expression level of rhEPO but their growth rate and survivability were astoundingly different in stressed condition (Fig.: 2A-D). Clone no. 2 grew at the lowest rate and Clone no. 3 grew at the highest rate, where the two other clones stayed in between. This data clearly showed the supremacy of the Clone no. 3 as the candidate for master cell bank (MCB) over the others.

**Figure 2:**
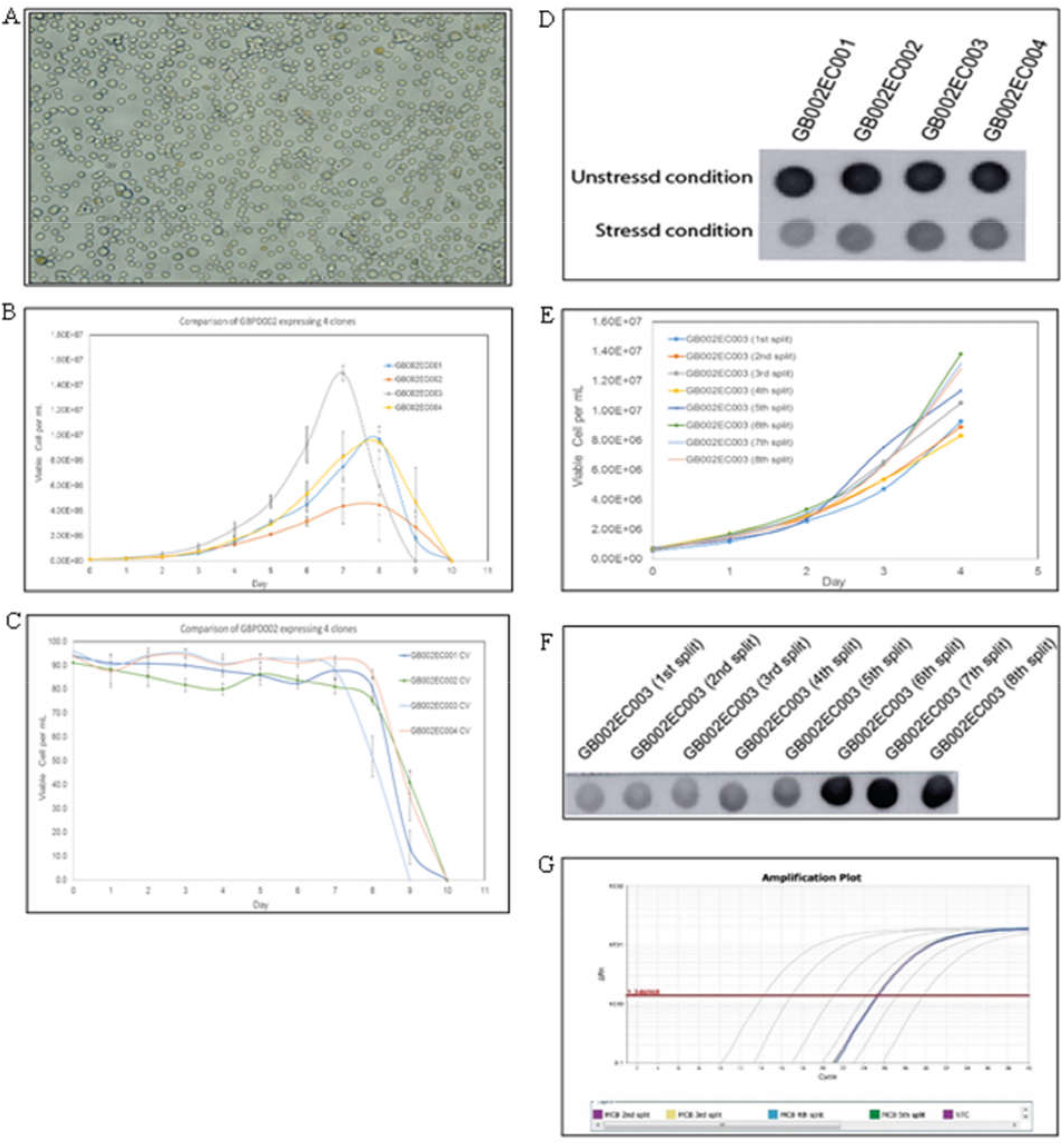
(A) Microscopic morphology of the clone, (B) Growth curve analysis of selected clones, (C) Viable percentage analysis of selected clones, (D) Expression level of rhEPO in stressed and unstressed condition (E) Growth curve analysis of MCB of first 8 splits (F) Protein expression analysis of MCB of first 8 splits (G) Copy number analysis amplification plot of the rhEPO gene in cells form first 5 splits.

### Master cell bank and working cell bank

The selected clone (clone no.: 3) were further adapted for 4 more weeks under constant shaking and serum-free media. The cells were split and media were replenished on every 4^th^ day. The cellular morphology was remained similar over the period but the growth kinetics and protein expression level of rhEPO were increased for the first 5 splits and then remain constant (Fig.: 2E-F). The final growth rate for the cells were found 0.67/day and the specific productivity of cells were found 5.89 pg/cell/hr. At this point, the copy number of the rhEPO gene in cells were determined as 4 copies/cell, which remained unchanged throughout the adaptation process (Fig.: 2G). These cells were declared as MCB, and WCB were prepared from this MCB. The MCB and WCB were stored in liquid nitrogen as well as in -150 ºC freezer and were subjected for characterization.

Cells from MCB were further maintained in culture for 50 passages with a split and media replenishment on every 4^th^ day. These media showed consistent expression of the rhEPO with same molecular migration pattern detected by Western blot (Fig.: 3A). Cells from all passages were stored as mentioned before. The cellular growth kinetics and the copy number of the rhEPO gene also were found consistent (Fig.: 3B-C). Separate batches of cell culture were run (1L batch size in wave bioreactor) using the thawed cells from the frozen stocks (MCB) of passage no. 1, 25, and 50. Cell growth kinetics, copy number of rhEPO, and the molecular identity of rhEPO protein by Western blot were found similar among these three batches (Fig.: 3D). The cells were found free from mycoplasma and bacterial contaminations (Fig.: 3E-F).

**Figure 3:**
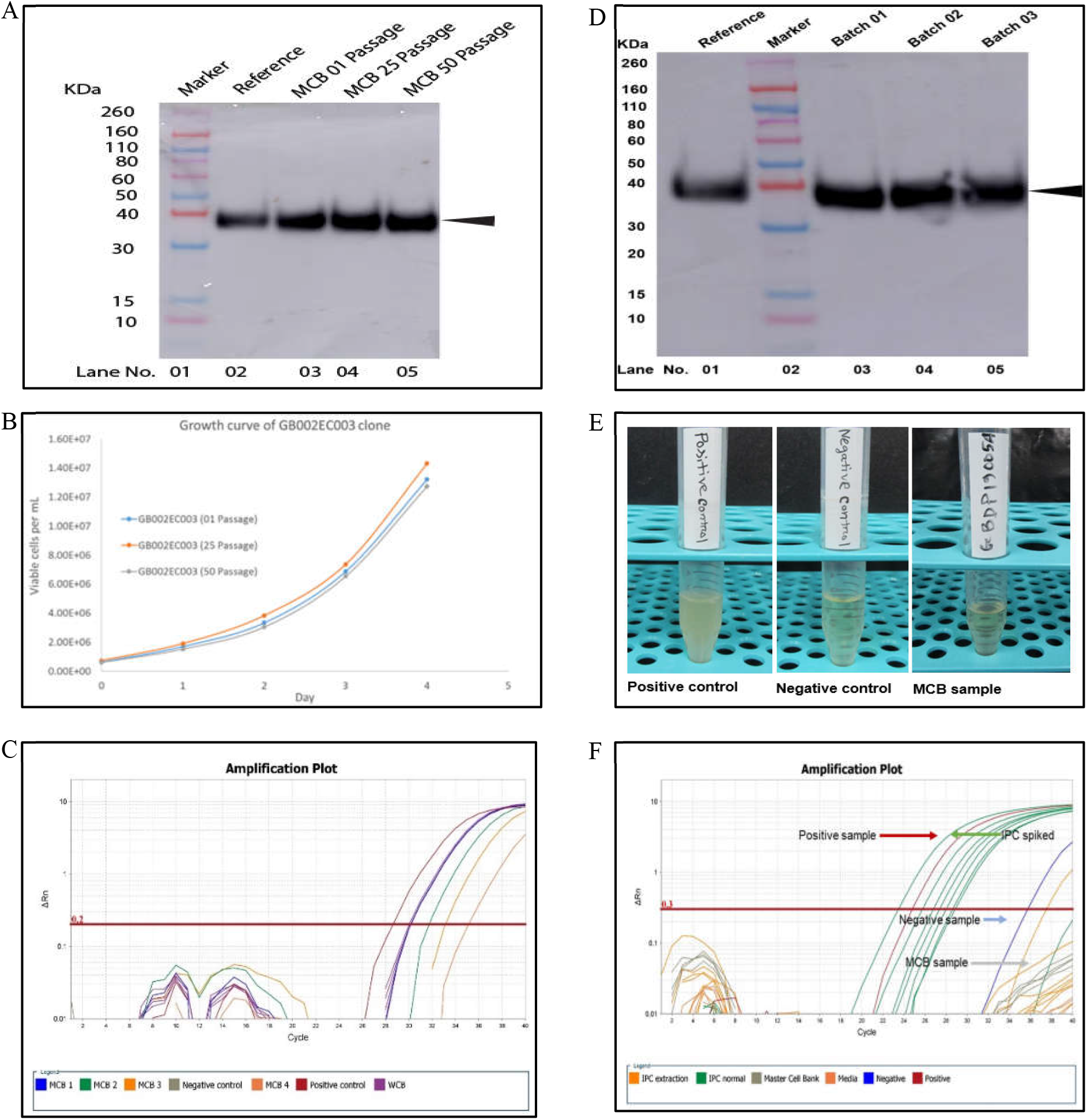
Cell line qualification of GBPD002. (A) Stability analysis for protein expression of MCB up to 50^th^ passages, (B) Growth kinetics of MCB for different passage, (C) Amplification plot of copy number analysis for rhEPO-insert of MCB, (D) Western blot analysis of 3 consecutives batches of MCB, (E) Microbial purity analysis of MCB by sterility testing, (G) Mycoplasma analysis of MCB.

### Manufacturing of rhEPO

Through a systematic approach, three consecutive batches were successfully performed in wave bioreactor using the WCB stock. During culture, maximum viable cell density (VCD) was above 18 million cells/ mL (total viable cell: 6.73 × 10^10 ± 5.85 × 10^8) at day 9 with viability 92% to 94%. Specific growth rate of cells was 0.0148 h-1 (day 2 to day 6), and decay rate were 0.0001 h-1 (day 6 to day 12) (Fig.: 4A-G). After harvesting the batches, the mass balance (amount of the total harvested media) for all these batches were found comparable (Batch 01: 4965 mL, Batch 02: 4980 mL, and Batch 03: 5010 mL). Relevant titer for the batches were found 1.23 g/L, 1.31 g/L, and 1.21 g/L, respectively for Batch 01, Batch 02, and Batch 03. The metabolite in the media were analyzed to check whether these batches might have any differences in relevant properties that may reflects differences in metabolic loads among the batches. We have tested ammonia, lactate, glucose, glutamate, and glutamine; all the trendlines were found in close proximity with no eccentric data point (Fig.: 5A-E). After the filtration, the appearance of the clarified samples was found brown with the absorbances of 1.79, 1.77 and 1.72, respectively at 600 nm for Batch 01, Batch 02, and Batch 03. The clarified sample did not produce any turbidity on standing at room temperature for 72 hours though the titer was reduced after 12 hours (Fig.: 5F), and suggested that the samples must be processed for the downstream within the next 12 hours.

**Figure 4:**
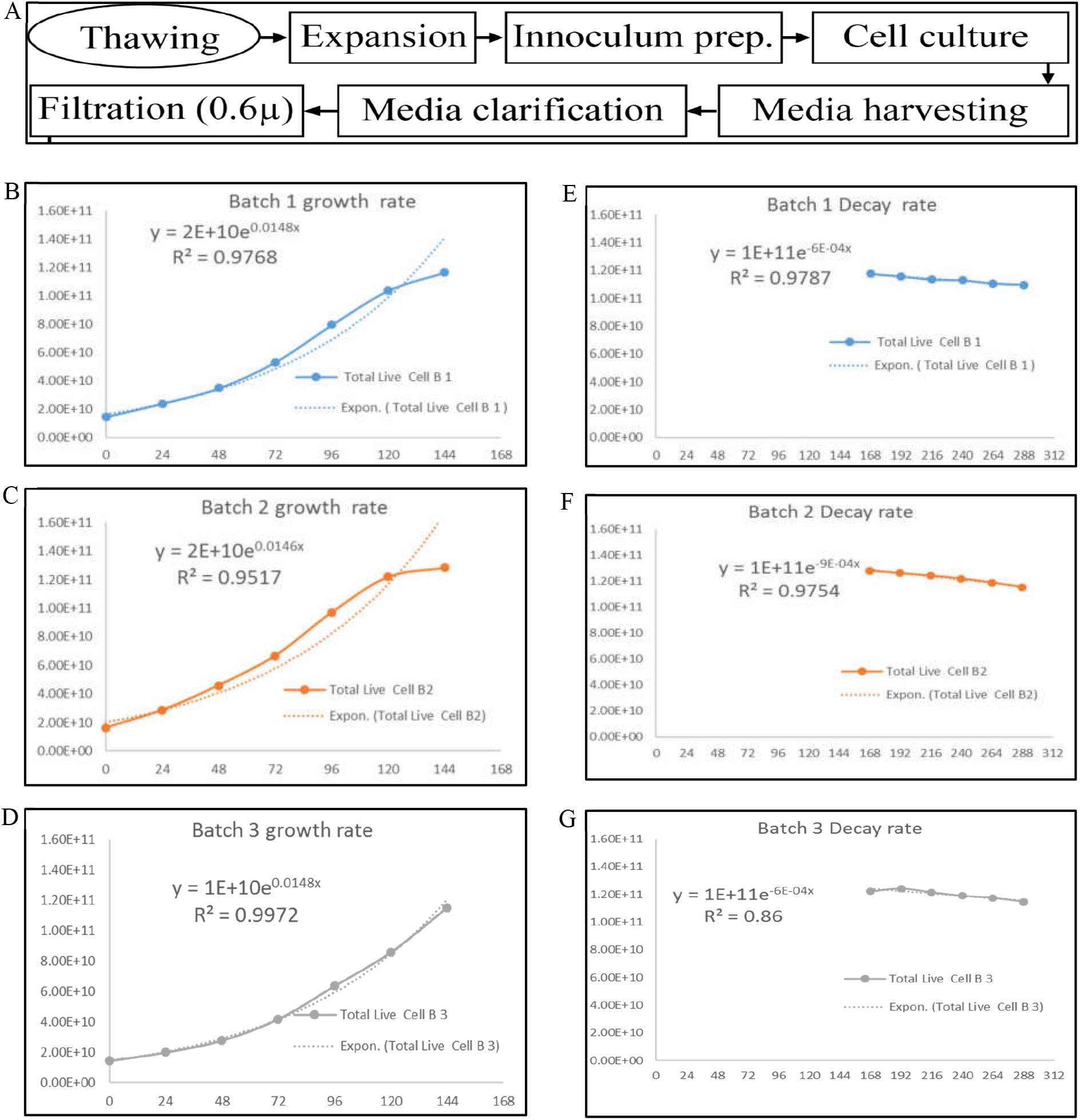
(A) Flow chart of Upstream process from thawing of WCB to harvest of cell culture media, (B) Growth rate of Batch 1 (C) Growth rate of Batch 2, (D) Growth rate of Batch 3, (E) Decay rate of Batch 1, (F) Decay rate of Batch 2, (G) Decay rate of Batch 3.

**Figure 5:**
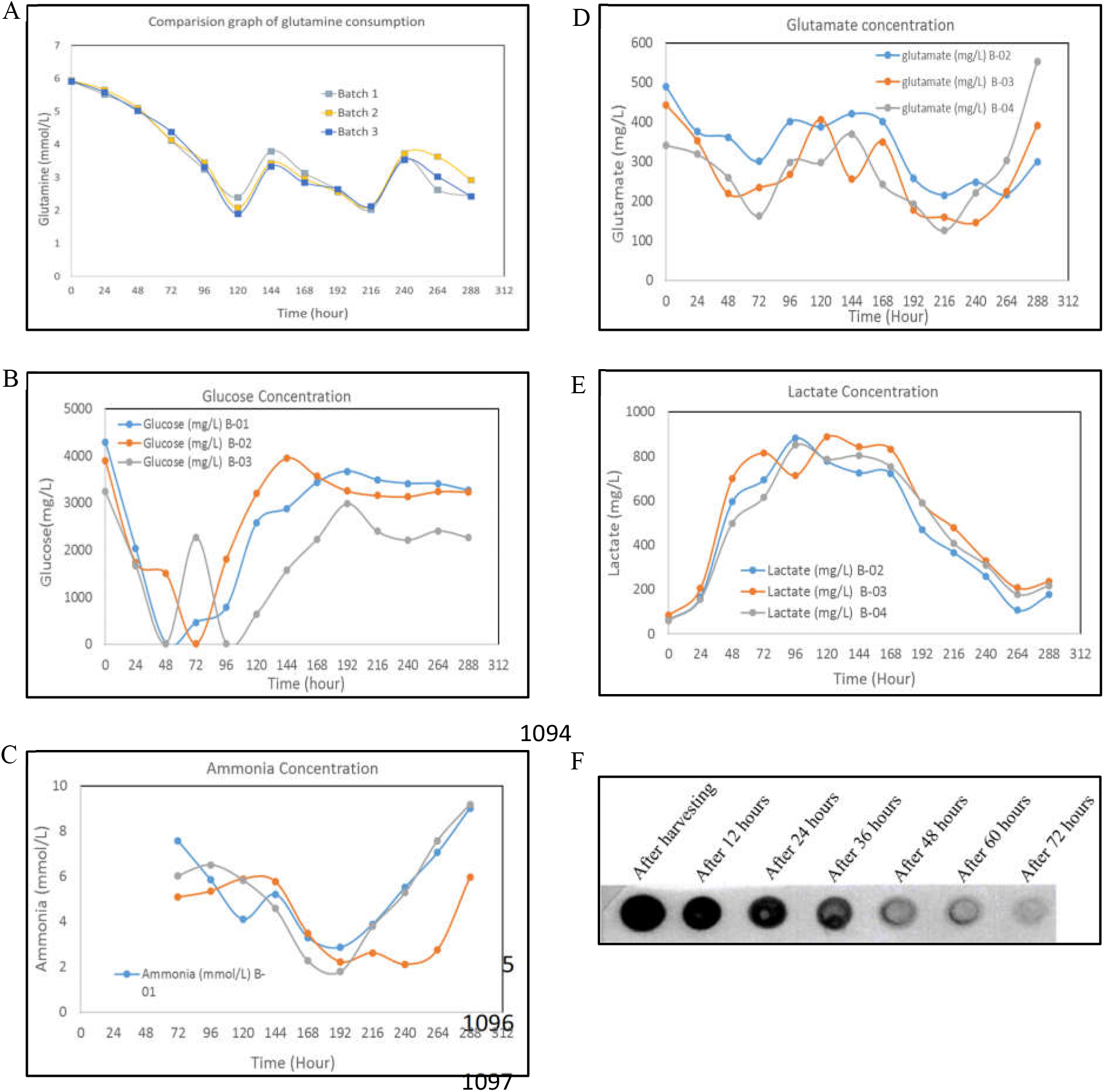
Metabolites analysis of batch 1 to 3. (A) Glutamine (B) Glucose (C) Ammonia, (D) Glutamate (E) Lactate (F) Sample holding stability after 12-hour interval at room temperature.

Accordingly, all batches were processed for the AFC to enrich the rhEPO and remove the majority of the unwanted materials as soon as possible after obtaining the media from the upstream processing. The rhEPO in final formulation were obtained by following the workflow shown in (Fig.: 6A.) At AFC unit process step, approximately 5.0 L of rhEPO-containing media sample was captured and eluted from resin; where sample pH, conductance, volume and UV280 were 7.34, 124.61 mS/cm, 5.0 L and 483,873 respectively. The eluate was buffer exchanged; where sample pH, conductance and volume were 7.12, 1.82 mS/cm, and 5.0 L, respectively. At AEX unit process step, approximately 1.5 L sample was eluted from resin; where sample pH, conductance, volume and UV280 were 7.05, 16.3 mS/cm, 1.5 L, and 305,150, respectively. At RPC unit process step, approximately 1.5 L sample was loaded and eluted from resin; where sample pH, conductance, volume, acetonitrile percentage and UV280 were 2.22, 2.79 mS/cm, 0.5 L, 41.6%, and 105,403, respectively. At VI-unit process step, 0.5 L RPC eluted sample was incubated for inactivation of virus or virus like particles; where incubation time and incubation temperature were 90 minutes and 22 ± 3 °C, respectively. At CEX unit process step, approximately 1.0 L sample was bound and eluted from resin; where sample pH, conductance, volume and UV280 were 6.98, 13.35 mS/cm, 0.84 L and 68,067, respectively. A set of relevant chromatograms are shown in (Fig.: 6B-E) The final yield for the three consecutive batches were found as follows: Batch 01: 0.724 gm, Batch 02: 0.728 gm, and Batch 03: 0.715 gm; the corresponding yield percentages were 13.30, 14.62, and 13.31, respectively.

**Figure 6:**
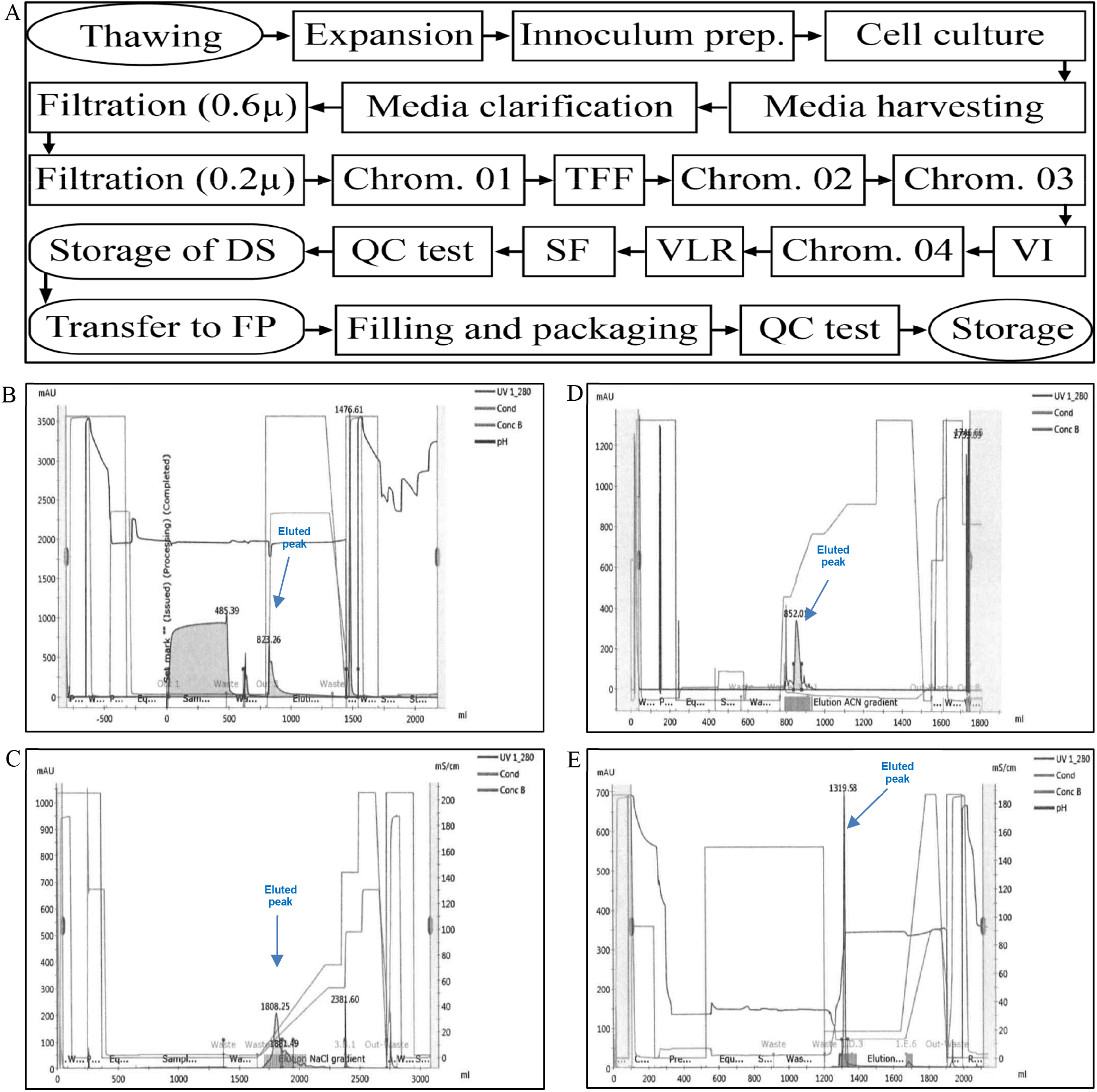
(A) Process flow of downstream process for GBPD002, (B) Representative chromatogram of affinity chromatography (AFC), (C) Representative chromatogram of anion exchange chromatography (AEX), (D) Representative chromatogram of reversed-phase chromatography (RPC), (E) Representative chromatogram of cation exchange chromatography (CEX) in the FPLC purification process.

### Qualification of the rhEPO formulation

The finished rhEPO was analyzed against the given specification (Table: 1). Since rhEPO is a parenteral preparation, therefore, the microbial sterility and endotoxin test were attempted first to conform with the specification and proceed forward with other critical quality attributes (CQA). No growth was found for microbial sterility testing, and the endotoxin values were found below 2.5 EU/mL for reference and GBPD002. To claim Eprex^®^ biosimilar, GBPD002 (Erythropoietin alfa) were analyzed by several identification tests. SDS PAGE and Western blot were performed for molecular mass and immunochemical detection analysis. High intensity similar banding pattern were found at near around 34 kDa for both reference and GBPD002 (Fig. 7A-B). two-dimensional gel electrophoresis showed well separated four isoform spots focused in the acidic region (pH 4.78-6.54) of the strips for both reference and GBPD002; though there were few other similar minor bands observed for both samples (Fig. 7C-D). Peptide mapping was performed by LC/MS-MS to obtain GBPD002 protein sequence coverage analysis. The results demonstrate 296 MS peaks, spanning 100% protein sequence coverage and 100% abundance (mol) (Fig. 7E). The pattern of MS peaks was very similar like those obtained from the reference product Eprex^®^ (Fig. 7F-G). Chromatographic pattern was also analyzed by reverse phase chromatography and results demonstrated similar retention (RT) time for both reference (9.550 min) and GBPD002 (9.558 min) (Fig. 7H), and suggested similar molecular identity for both proteins. Deglycosylation of N-linked sugars reduced the protein to 18 kD from the original protein of ∼37kD; the reference protein also showed similar protein bands on deglycosylation as well as for full form (Fig. 8A-B).

**Table 1:** Head-to-Head analysis of Eprex^®^ and GBPD002.

| Test Parameters | Eprex<sup>®</sup> | GBPD002 |
| --- | --- | --- |
| **Structural Analysis** |  |  |
| Molecular mass and immunochemical properties analysis by SDS PAGE and Western Blot | High intensity similar banding pattern were found at near around 37 kDa | High intensity similar banding pattern were found at near around 37 kDa |
| Peptide Mapping by LC-MS/MS | 100% protein sequence coverage and 100% abundance (mol) | 100% protein sequence coverage and 100% abundance (mol) |
| Isoform pattern analysis by 2D gel electrophoresis (2D GE) | Well separated, four isoform spots focused in the acidic region (pH 4.78-6.54) of the strip | Well separated, four isoform spots focused in the acidic region (pH 4.7-6.5) of the strip |
| Chromatographic pattern analysis by HPLC | Chromatogram pattern and retention time are similar<br>Retention time (RT) time: 9.550 | Chromatogram pattern and retention time are similar<br>Retention time (RT) time: 9.558 |
| **Impurity Analysis** |  |  |
| Aggregation pattern analysis by size-exclusion chromatography (SEC) | Single similar type of peak was found | Single similar type of peak was found |
| Particle size analysis by Malvern zeta sizer | No aggregated and degraded peak was found<br>Z-average was found 11.50 ± 0.81 nm with the PDI values 0.294 | No aggregated and degraded peak was found<br>Z-average was found 11.49 ± 0.28 nm with the PDI values 0.218 |
| **Functional Analysis** |  |  |
| Kinetics analysis by surface plasmon resonance (SPR) | The association rate ($k_a$) was $2.916E^5$ and the dissociation rate was ($k_d$) $2.538E^{-8}$ | The association rate ($k_a$) was $3.124E^5$ and the dissociation rate was ($k_d$) $2.838E^{-8}$ |
| Cell culture-based assay | High cell proliferation was found at dose concentration 2 ng/mL | High cell proliferation was found at dose concentration 2 ng/mL |
| In-vivo study by mice model | Reticulocytes (Ret) were counted as the primary marker.<br>Ret:<br>Placebo: $3.5 \pm 0.54$ %.<br>Control: $3.9 \pm 0.85$ %.<br>Treatment-I (20 IU/ mL: $10.0 \pm 1.17$ %.<br>Treatment-II (40 IU/ mL: $12.5 \pm 0.60$ %.<br>Treatment-III (40 IU/ mL: $13.5 \pm 0.54$ %. | Reticulocytes (Ret) were counted as the primary marker.<br>Ret:<br>Placebo: $3.5 \pm 0.54$ %.<br>Control: $3.9 \pm 0.85$ %.<br>Treatment-I (20 IU/ mL: $10.3 \pm 0.45$ %.<br>Treatment-II (40 IU/ mL: $12.9 \pm 0.53$ %.<br>Treatment-III (40 IU/ mL: $13.9 \pm 0.45$ %. |
| Assay by reverse phase chromatography | Chromatographic pattern complies the testing acceptance criteria | Chromatographic pattern complies the testing acceptance criteria ($103.0 \pm 2.0\%$) for 3 consecutive batches were $103.0 \pm 2.0\%$. |

**Figure 7:**
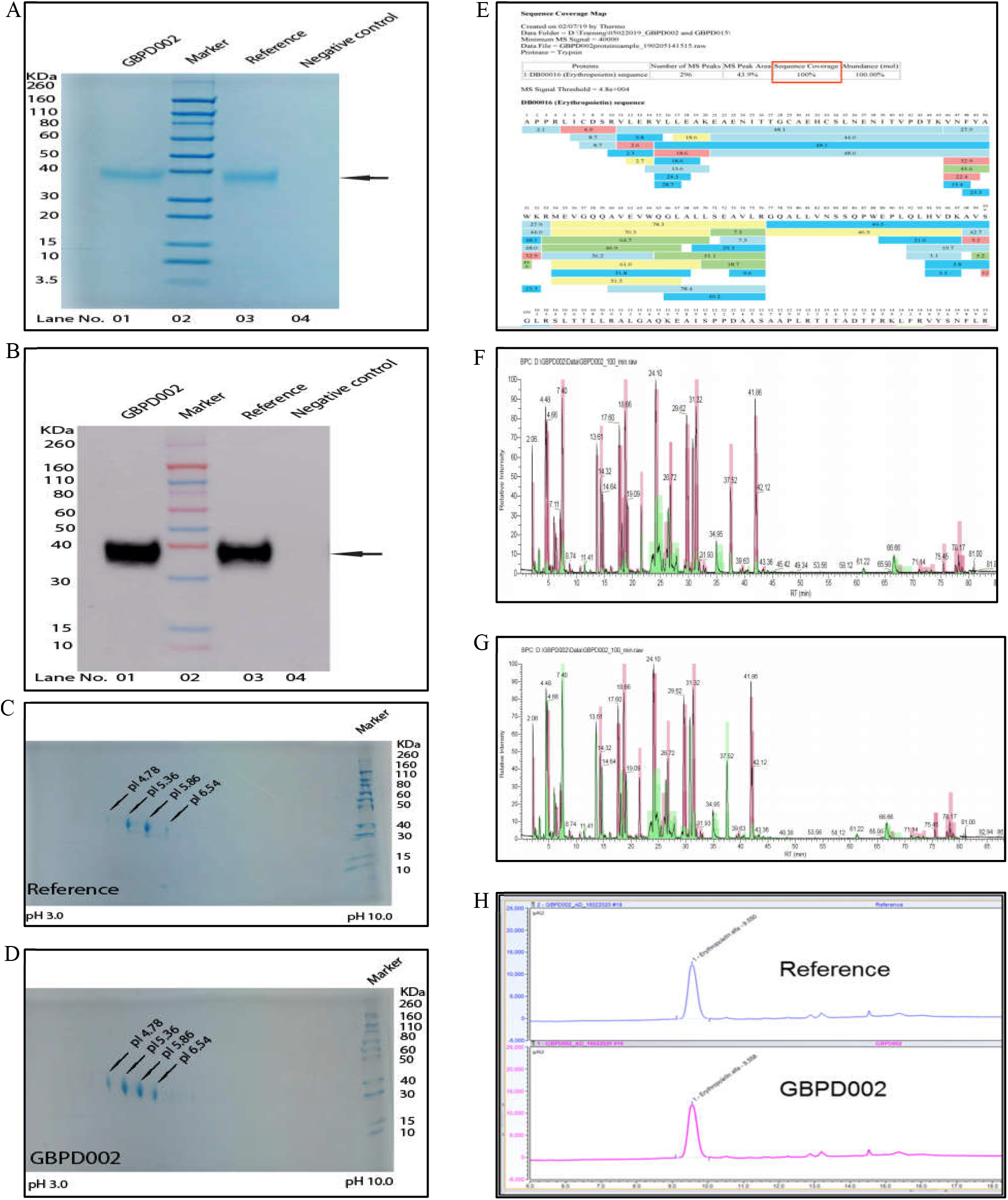
Comparative analysis for identification test between reference product Eprex^®^ and GBPD002. (A) SDS PAGE analysis, (B) Western Blot analysis, (C) Isoform pattern analysis of Eprex^®^, (D) Isoform pattern analysis of GBPD002, (E) Peptide mapping (amino acid sequence) of GBPD002, (F) Peptide mapping chromatogram (relative intensity) of Eprex^®^, (G) Peptide mapping chromatogram (relative intensity) of GBPD002, (H) Comparative study for chromatographic pattern analysis by Reverse-phase chromatography (RPC).

**Figure 8:**
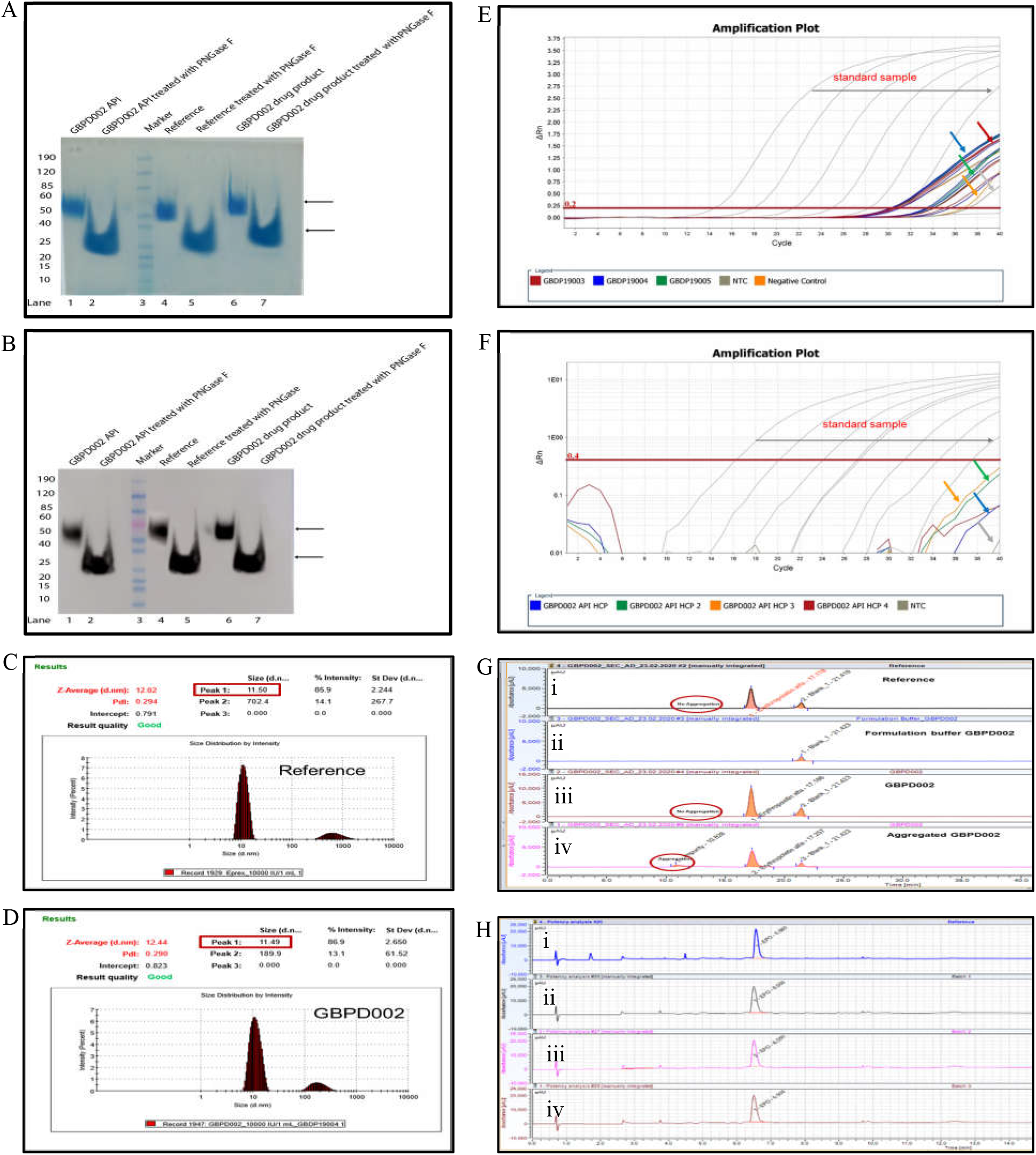
Comparative analysis for glycosylation pattern, process- and product-related impurities between Eprex^®^ and GBPD002. (A) Glycosylation pattern analysis by SDS PAGE, (B) Glycosylation pattern analysis by western blot, (D) Particle size analysis by Malvern zeta sizer of Eprex^®^, (E) Particle size analysis by Malvern zeta sizer of GBPD002, (E) Host cell DNA analysis (process-related impurities), (F) Host cell protein analysis (process-related impurities), (G) Aggregation pattern analysis (product-related impurities) by size-exclusion chromatography (SEC), (H) Potency and quantity analysis by Reverse-phase chromatography (RPC).

Macromolecular aggregation and particle size distribution were analyzed. For Particle size analysis, Z-average were found 11.50 ± 0.81 nm and 11.49 ± 0.28 nm with the PDI values 0.294 and 0.218 for reference and GBPD002, respectively (Fig.8C-D). Host cell DNA and host cell protein (HCP) were analyzed by real-time quantitative PCR (RT-qPCR). For Host cell DNA, three consecutive batches were analyzed and results demonstrated <10 ng/sample for all 3 batches (Fig. 8E). Three consecutive batches were analyzed and HCP concentrations were found below the detection limit (Fig. 8F), which has indicated that the rhEPO preparations were having non-detectable HCP. SEC revealed a single peak in chromatogram for the reference and GBPD002 product (Fig.8 Gi). There was a noticeable secondary peak observed for both samples, which can be attributed to the buffer components (Fig. 8 Gii). No other aggregated and/or degraded peak was found for any samples (Fig. 8 Giii). RPC analyses revealed the assay for the GBPD002 preparations for 3 consecutive batches were 103.0 ± 2.0% in comparison with the reference (Fig. 8H).

### Biofunctional activity

Comparative functional study was performed by receptor binding assay, *in vitro* cell culture assay, and *in vivo* bioassay between reference and GBPD002 to prove the similar bio-functionality between the reference and GBPD002. Receptor binding assay against immobilized EPO receptor revealed similar types of interaction kinetics for both reference and GBPD002. The association rate (ka) was found 2.916E5 and 3.124E5 for reference and GBPD002, respectively; whereas the dissociation rate was found (kd) were 2.538E-8 and 2.838E-8 for reference and GBPD002, respectively (Fig. 9A-B). The experiment U-value was 9.

**Figure 9:**
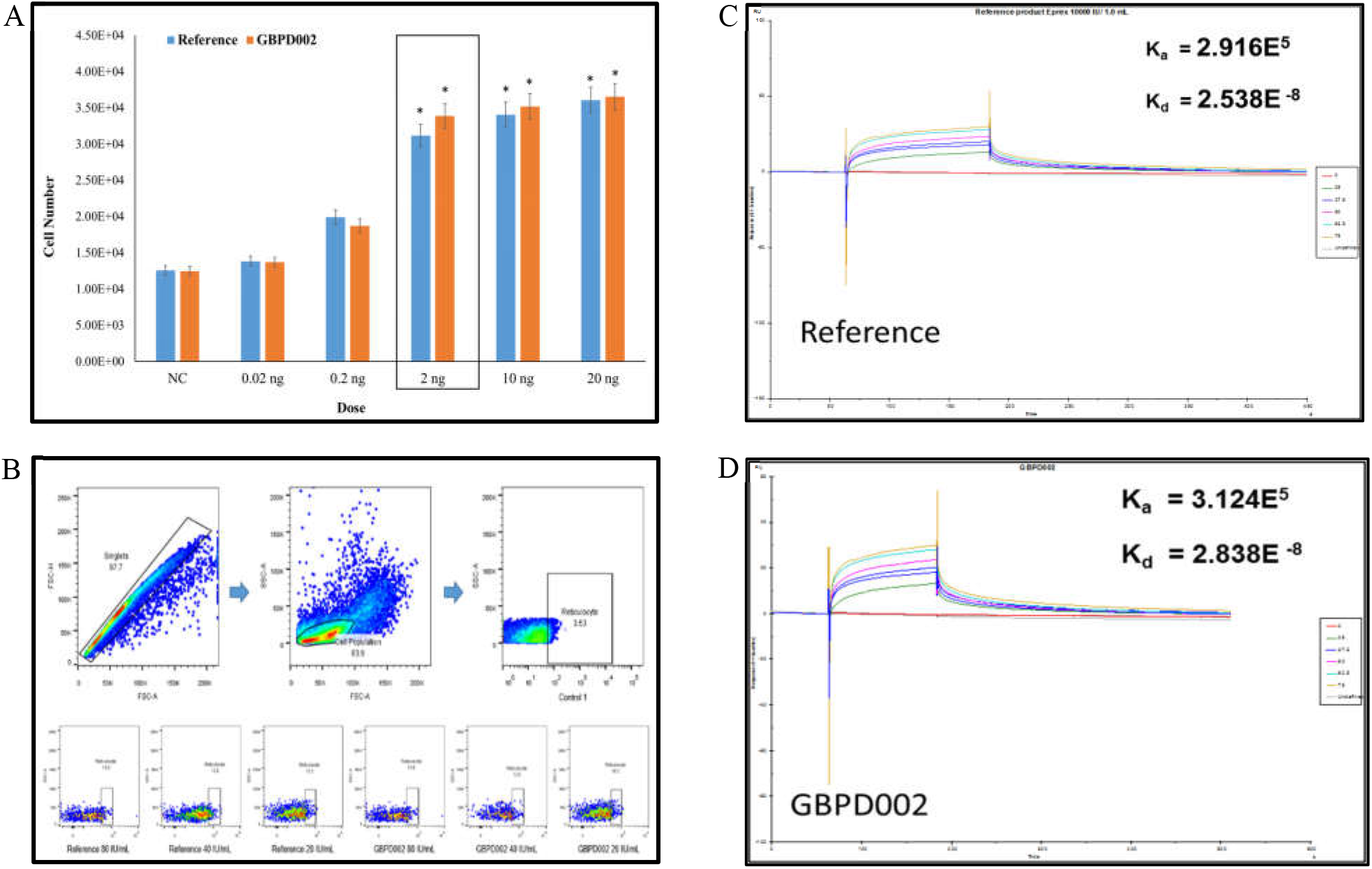
Comparative analysis of bio-functionality between Eprex^®^ and GBPD002. (A) Graphical presentation of cell culture-based assay, (B) *In-vivo* study in mice model, (C) Analysis of receptor binding kinetics by SPR of Eprex^®^, (D) Analysis of receptor binding kinetics by SPR of GBPD002.

The rhEPO-induced proliferation data of TF-1 cell clearly revealed that the GBPD002 and reference products responded similarly. At a 2 ng/ml and higher concentration, both of the rhEPO formulations assisted the growth of experimental cells by 3 times of the originally seeded cell numbers (Figure 9C). The mock-controlled cells, on the other hand, were reduced to half by number from its original seeding population (10^5^/well). This data has clearly established that the rhEPO preparations GBPD002 and Eprex^®^ are similarly active.

For *in vivo* bioassay, mice reticulocytes (Ret) were counted as the primary marker. The mean values of Ret were found 3.5 ± 0.54 % and 3.9 ± 0.85 % for placebo and control group, respectively. As expected, for treatment-I (20 IU/ mL), the mean values of Ret were found increased to 10.0 ± 1.17 % and 10.3 ± 0.45 % for reference and GBPD002, respectively. For treatment-II (40 IU/ mL), the mean values of Ret were found 12.5 ± 0.60% for reference and 12.9 ± 0.53 % for GBPD002, respectively. For treatment-III (80 IU/ mL), the mean value of Ret was found (13.5 ± 0.54) and (13.9 ± 0.45) % for reference and GBPD002, respectively (Fig. 9D).

## 4. DISCUSSION

Two rhEPO products were compared to determine biosimilarity. These were the reference product Eprex^®^ (International Nonproprietary Name: epoetin alfa) and a candidate biosimilar of rhEPO designated as GBPD002. Both were prepared using rDNA technology, expressed in suspension culture of CHO cells, and purified using a series of validated unit processes. We have chosen CHO cells because the reference product is also CHO cell derived. Furthermore, it has been shown that CHO cell has similar level of glycosylation-related transcripts to human cells [35], which may help to obtain the expressed protein with desired post-translational modification (PTM). CHO cells can also evade infection of human virus through minimizing the expression of many important viral entry gene that improves final product safety [35].

Several techniques have been in use for development and isolation of stable mammalian cell lines, e.g., FACS, automated colony picker, serial infinite dilution etc. FACS needs addition of foreign biological reagent like antibody as well as application of photo-beam, which may harm cell viability and cellular integrity [36]. FACS and automated clone picker (such as Clonepix and CellCelector) also need highly expensive instrumentation. On the contrary, classical limiting dilution technique, though time consuming but does not require expensive instrumentation, is the best method to harvest single clones satisfying the regulatory requirements [37]. Therefore, this classical method for clonal isolation of desired cells was followed in the study. A common practice is to use antibiotic challenge at a suitable concentration to select the stably-transfected cells. Some low-expressing or unstably-transfected or non-transfected cells also can survive in this process, albeit with a lower frequency. In this study we have used highest possible concentration of antibiotic challenge to identify the highest-expressing cells with only a few cells surviving on challenge. Such high level of selection challenge was also applied with Methotrexate (MTX) selection for dihydrofolate reductase (DHFR) compromised cells resulting selection of high-level protein expressing clones [38-40].

The cell growth rate was compromised during the high-concentration antibiotic challenge, which is a common phenomenon for such protocol [41] however, during the subsequent adaptation process, the growth rates for the clones were revived. The isolated clones usually go for evaluation of growth rate and the specific productivity per cell to identify the best desired clone for further development to MCB and WCB. We have applied a new screening technique where the clones whose growth kinetics and specific expressions were comparable were subjected for a capacity test in a fixed media depriving fresh supplements and with accumulating load of metabolites. This stress-test revealed a superior clone over the others, which performed exceptionally in batch format to produce high level of expression for rhEPO (1.24±0.16 g/L). To the best of our knowledge, this is the highest level of titer for rhEPO expression using a standard fed-batch wave bioreactor system. Searching of available information in web space revealed the highest level of rhEPO titer was ∼696 mg/L for a transient expression system [42].

In 2013, the World Health Organization (WHO) published a Technical Report Series (TRS 978: Recommendations for the Evaluation of Animal Cell Cultures as Substrates for the Manufacture of Biological Medicinal Products and for the Characterization of Cell Banks) that states, “For proteins derived from transfection with recombinant plasmid DNA technology, a single, fully documented round of cloning is sufficient provided product homogeneity and consistent characteristics are demonstrated throughout the production process and within a defined cell age beyond the production process”. Accordingly, we have successfully completed the isolation of single clones and expanded them, and finally characterized them to qualify as the MCB. Cell cultures are maintained for not more than 60 days for a N-6 systems (generally, each step may consume 7 days each for up to N-5 steps (5×7=35 days), and the 6^th^ step may consume 18 days, total 35+18=53 days) in manufacturing train of a commercial batch (2000 L) in fed-batch process. Continuous batch run time in perfusion mode also generally does not run for more than 6 months (180 days). Here, we have characterized the MCB for around 200 days, which can be used for manufacturing of rhEPO either in fed-batch or continuous mode without compromising yield and product quality.

Genomic plasticity is inherent to any immortalized mammalian cell lines [43-45]. Recently, CHO genome drafts revealed significant chromosomal heterogeneity among different CHO cell lineages and genomic landscapes [35, 46]. Therefore, clonally isolated production cells will undergo some genetic heterogeneity over the age, though different clones will withstand original property for different ages[44, 45, 47-49]. We presumed that the exceptional stability of our MCB for product substrate is due to the acquired fitness of the clone associated with the sequence of multiple stringent screening methods during clonal isolation. This notion is supported by a recent study, which has reported that a secondary screening of an established MCB produced high-titer stable clones without affecting the quality attributes of the desired product [50].

A working guideline has been suggested for setting the specification for the qualification of biological drugs [51]. Further, through a case study, the quality assessment framework has been suggested for rhEPO biosimilar characterization by regulatory experts [52]. We have followed these suggestions and performed the characterization with the scientifically reasonable experimental methods. One of the first critical quality attributes was the cell substrate. As mentioned in other reference, [53] ICH guideline Q5D Section 2.1.3 states that “For recombinant products, the cell substrate is the transfected cell containing the desired sequences, which has been cloned from a single cell progenitor”, we have deduced the gDNA and cDNA sequences for the rhEPO insertion in the MCB, which was completely matched with the reference sequence. Amino acid sequencing form the formulated rhEPO sample also showed 100% match with the protein sequence in data bank. These data confirmed the cell substrate conformity for the MCB and WCB for rhEPO production.

The identification of the rhEPO was done with orthogonal methods SDS-PAGE, Western blot, dot blot, Elisa, as well as UHPLC. All methods explicitly confirmed the similar molecular pattern and immunochemical properties for GBPD002 and Eprex®. The biosimilar comparison included peptide mapping and mass spectrometry procedures to asses amino acid sequence and mass spectra, which were found highly similar with a few minor variations. Sialic acid content is very important components among the glycosylation matrix, and is directly connected to the half-life of the rhEPO for in vivo functionality. Similar charge variant profiles of GBPD002 and Eprex^®^ suggested likely similar silic acid contents for the both rhEPO preparations. Since both products likely have similar sialic acid content, therefore, such minor variations can be attributed to differences in sugar residues related to glycosylation. PNGase F digestion of protein followed by molecular mobility detection has been used for analysis of glycosylation pattern of proteins [54]. Accordingly, we have also applied the same method for determining comparability for glycosylation profile of GBPD002 and Eprex®. The molecular mobility and immunodetection patterns of full-form and deglycosylated form were found highly similar for GBPD002 and Eprex® thereby suggesting that the differences in glycosylation between the two products are minimum, if there is any.

Glycosylation profile has been found important for protein folding that affects protein stability and receptor interaction. However, for rhEPO exact glycosylation between two products seems not rational as discussed below: (1) Non-glycosylated Escherichia coli-expressed rEPO have demonstrated that it is well folded and possess the same *in vitro* biological activity as the CHO cell-derived glycosylated rEPO [55, 56]. (2) Multiple different EPO-glycoforms were identified in circulating systems of human [57]. (3) The glycosylation profile of human serum EPO is remarkably different from the rhEPO [58]. (4) There were no significant differences found for *in vivo* biofunctionality for several approved rhEPO preparations (12 different sources) despite having differences in glycosylation profile [59]. (5) Several clinical studies did not find any significant differences between pharmacokinetic and pharmacodynamic properties for different rhEPO [60-66]. (6) Confident glycoform analysis is a very critical process, and multiple parallel methods are preferred for obtaining confident PTM profiles using mass spectrometry analysis [67, 68], which might not be supportive for cost-benefit perspective. (7) Further, it has been suggested that the separation of rEPO into pure glycoforms is not feasible even with the most advanced methods due to the heterogeneity of the three N-glycans [69]. (8) Minor differences in glycosylation may not be affecting the final product quality significantly. In reality, such type of minor variations are not infrequent for EMA and US FDA approved biosimilars [70]. A recent study reported the batch-to-batch variability in quality parameters of marketed rhEPO reference medicines, Eprex®/Erypo® (Janssen-Cilag, High Wycombe, UK) and NeoRecormon® (epoetin beta; Roche Registration Limited, Welwyn Garden City, UK), and two biosimilars, Binocrit® (Sandoz GmbH, Kundl, Austria) and Retacrit® (Hospira UK Limited, Maidenhead, UK), and found batch-batch minor variability for experimental products [71]. Therefore, it has been suggested that, it is not essential to have identical glycosylation pattern for two molecules to be considered biosimilar though the biofunctionality must conform to offset the residual uncertainty [32, 52].

To offset the residual uncertainty, we have performed the receptor binding assay, and bio-functional assay in mammalian cell as well as in mice model. The receptor binding assay, and the *in vitro* and *in vivo* biofunctionality assays clearly demonstrated very similar receptor binding affinity and biological activities for GBPD002 and Eprex^®^. EPO-specific antidrug antibodies (ADAs) are generated *via* immunogenicity of an administered rhEPO preparation, and they likely cross-react with endogenous EPO in human. Aggregation present in rhEPO preparation were held responsible for the ADA effect. These ADAs, in severe cases, lead to clinically relevant autoimmunity known as pure red cell aplasia (PRCA) [72-74]. Three orthologous analysis procedures, *viz*., Western blot, dynamic light scattering and SEC-HPLC were applied for detecting any high molecular aggregates in our final products, and the results confirmed that GBPD002 did not contain any such macromolecular entity. Together, all the results clearly demonstrated the biosimilarity of GBPD002 and Eprex^®^.

## 5. CONCLUSION

The study showed systematic development of a rhEPO biosimilar from the DNA preparation to the formulation following regulatory guidelines. Higher yield of the relevant process can help reducing the product cost and may socioeconomically support the global communities who are in need of rhEPO at a comparatively cheaper price. The physicochemical and biofunctionality results suggested the comparable biosimilarity between GBPD002 and Eprex^®^. Nevertheless, the complete similarity for biofunctionality should be investigated in a clinical study with ‘head-to-head’ comparability between GBPD002 and the reference and should include appropriate endpoints, *e*.*g*., hematocrit and hemoglobin. Such a study would reveal clinically noteworthy variance in the *in vivo* quality attributes for the rhEPO preparations.

## Author contribution

Kakon Nag and Naznin Sultana conceptualized the project. Md. Jikrul Islam, Md. Maksusdur Rahman Khan, Md. Mashfiqur Rahman Chowdhury, Md. Enamul Haq Sarker and Samir Kumar designed experiments and analyzed data. Md. Maksusdur Rahman Khan, Rony Roy and Md. Shamsul Kaunain Oli contributed to cell line development, and Md. Mashfiqur Rahman Chowdhury contributed to cell bank development and scale-up of the cell culture. Md. Enamul Haq Sarker, Samir Kumar, Sourav Chakraborty and Habiba Khan performed downstream process development and manufacturing steps. Md. Jikrul Islam and Ratan Roy performed quality control experiments. Kakon Nag, Naznin Sultana, Md. Jikrul Islam, Md. Maksusdur Rahman Khan, Md. Mashfiqur Rahman Chowdhury, Md. Enamul Haq Sarker and Samir Kumar wrote the manuscript. Bipul Kumar Biswas, Md. Emrul Hasan Bappi and Mohammad Mohiuddin assured the quality management system of relevant activities.

## Institutional review board statement

Not applicable.

## Informed consent statement

Not applicable.

## Data availability statement

The data that support the findings of this study are available within the article and its Supplementary document file, or are available from the corresponding author upon reasonable request.

## 6.0 ACKNOWLEDGEMENTS

The study was funded by Globe Biotech Limited. We thank Md. Harunur Rashid, the chairman of Globe Pharmaceuticals Group of Companies, Ahmed Hossain, Md. Mamunur Rashid and Md. Shahiduddin Alamgir, Abdullah Al Rashid, the directors of Globe Pharmaceuticals Group of Companies for their continuous support and encouragement. We also thank Md. Raihanul Hoque, Dibakor Paul, Zahir Uddin Babor, Mithun Kumar Nag, Alok Sutradhar, G.M. Sajib Hasan, Biplob Biswas, and Mijanur Rahman for their support for facility and information management system.

